# Toward a 3D model of phyllotaxis based on a biochemically plausible auxin-transport mechanism

**DOI:** 10.1101/398628

**Authors:** Félix P. Hartmann, Pierre Barbier de Reuille, Cris Kuhlemeier

## Abstract

Polar auxin transport lies at the core of many self-organizing phenomena sustaining continuous plant organogenesis. In angiosperms, the shoot apical meristem is a potentially unique system in which the two main modes of auxin-driven patterning— convergence and canalization—co-occur in a coordinated manner and in a fully three-dimensional geometry. In the epidermal layer, convergence points form, from which auxin is canalized towards inner tissue. Each of these two patterning processes has been extensively investigated separately, but the integration of both in the shoot apical meristem remains poorly understood. We present here a first attempt of a three-dimensional model of auxin-driven patterning during phyllotaxis. We base our simulations on a biochemically plausible mechanism of auxin transport proposed by Cieslak et al. (2015) which generates both convergence and canalization patterns. We are able to reproduce most of the dynamics of PIN1 polarization in the meristem, and we explore how the epidermal and inner cell layers act in concert during phyllotaxis. In addition, we discuss the mechanism by which initiating veins connect to the already existing vascular system.

**Author summary:** The regularity of leaf arrangement around stems has long puzzled scientists. The key role played by the plant hormone auxin is now well established. On the surface of the tissue responsible for leaf formation, auxin accumulates at several points, from which new leaves eventually emerge. Auxin also guides the progression of new veins from the nascent leaves to the vascular system of the plant. Models of auxin transport have been developed to explain either auxin accumulation or auxin-driven venation. We propose the first three-dimensional model embracing both phenomena using a unifying mechanism of auxin transport. This integrative approach allows an assessment of our present knowledge on how auxin contributes to the early development of leaves. Our model reproduces many observations of auxin dynamics. It highlights how the inner and epidermal tissues act together to position new leaves. We also show that an additional, yet unknown, mechanism is required to attract new developing veins towards the main vasculature of the plant.

## Introduction

In plants, most developmental processes are driven by the spatiotemporal distribution of the growth regulator auxin. The versatility of the morphogenetic role played by auxin relies on its self-regulated polar transport, in which auxin transport feedbacks on auxin efflux carriers (PIN proteins). This process can lead to various distribution patterns, depending on the specific geometry of the organ considered. Although auxin transport and patterning occur fundamentally in three-dimensional tissues, they are usually explored in one- or two-dimensional models. These restrictions are commonly justified by geometrical considerations. For instance, models of phyllotactic patterning in the shoot apical meristem (SAM) of *Arabidopsis* assume that the formation of convergence points of PIN1 polarization at primordia takes place in the single epidermal cell layer L1 [1-4], based on observations by Reinhardt et al. [5]. However, there is evidence that inner tissues may also be involved in positioning primordia [6,7].

Bayer et al. [8] proposed an auxin transport model integrating both the formation of convergence points in the L1 and the patterning of vascular strands in the subepidermal layers. Their model is implemented on a 2D cellular template representing a longitudinal section through the meristem. Thus, primordia positioning and midvein development are simulated in, respectively, one and two dimensions. These restrictions in dimensionality limit the range of potential behaviors displayed by the model, and therefore its capacity to integrate and assess current knowledge on phyllotaxis. Although fully three-dimensional models would be a significant step forward, they still present technical challenges [9]. In a notable effort in this direction, Gruel et al. [10] modeled gene expression dynamics and cell-to-cell diffusion of signals on a 3D cellular template to explore positioning and maintenance of the stem cell niche in the meristem. Modeling auxin polar transport is arguably more challenging, since it requires in addition a description of PIN exo- and endocytosis between cytoplasm and membranes, and, ideally, auxin diffusion in the apoplast. Several technical difficulties pertaining to the modeling of 3D plant tissues could be alleviated by using the topological notion of cell complexes [11]. An implementation of this paradigm has been utilized by Yoshida et al. [12] to investigate the control of division orientation in early *Arabidopsis* embryogenesis. Cell complexes are likely to turn out even more useful when fluxes between tissue components (cytoplasm, membranes, cell walls) are involved.

Another defying peculiarity of the SAM is the co-occurrence of convergence and canalization at midvein initiation. In the L1, PIN1 polarize towards convergence points, where auxin accumulates. From these points, strands of cells with high PIN1 expression extend into the subepidermal layers, with PINs displaying a canalization pattern. This co-occurrence suggests that these two modes of auxin transport regulation do not rely on completely different mechanisms, but instead relate somehow to each other.

Convergence and canalization are commonly conceptualized using two distinct polarization models, respectively referred as “up-the-gradient” or “with-the-flux”. There has been interest to go beyond this dichotomy. Some models attempted to explain phyllotaxis with either purely up-the-gradient [13] or with-the-flux [4] polarization, but they lack biological plausibility or contradict experimental data [14]. Following another approach, Bayer et al. [8] put forward the concept of “dual polarization”, whereby both polarization mechanisms operate concurrently, with a continuous transition from up-the-gradient to with-the-flux polarization depending on local auxin concentration. Although the dual-polarization model displays good agreement with observations, it remains difficult to explain how two qualitatively different mechanisms of PIN1 localization can coexist within the same cells. Shifting attention from A*rabidopsis* to *Brachypodium*, O’Connor et al. [15] took advantage of the existence of the PIN1 duplicate sister-of-PIN1 (SoPIN1) proteins in grasses by attributing different roles to the two proteins. However, no such duplication exists in A*rabidopsis*, where both convergence and canalization are apparently performed by PIN1 protein alone.

Moreover, there are fundamental problems with most polarization models [14]. In with-the-flux polarization, it seems unrealistic that cells directly sense net fluxes of auxin through their membranes. Regarding up-the-gradient polarization, it is not clear how a cell could react to the auxin concentration in their neighbors. Several alternative mechanisms have been proposed. Wabnik et al. [16] assumed that auxin gradients in the apoplast are informative enough to drive polarization. Although their model is capable of transitioning between up-the-gradient and with-the-flux regimes, and might thus also account for phyllotaxis, it is yet to be determined whether significant auxin gradients can form in the very narrow spaces between meristematic cells. Abley et al. [17] proposed a model for tissue polarity based on cell-cell coupling through a diffusive mediating molecule. They used it to reproduce several polarization behaviors [18], but phyllotactic patterns and midveins initiation seem out of reach. A further strategy has been to incorporate biomechanical factors [19], which can generate a whorled pattern of auxin maxima, but not the range of polarizations observed in phyllotaxis.

In all up-the-gradient models, PIN polarization is assumed to be fast compared to the production and turnover of PINs, as well as to changes in cellular auxin concentration, so that PIN concentrations at membranes are set to their steady-state values [9]. Since the cycling rates of PINs are completely unknown [14], this assumption may not be valid. If there is indeed significant latency between changes in auxin concentration and PIN polarization, up-the-gradient models could fail to capture some aspects of convergence point formation.

In an attempt to explain how cells can measure the direction and magnitude of auxin fluxes, Coen et al. [20] hypothesized the existence of “tally molecules” produced or consumed at the membrane when auxin enters/exits the cell. The concentration of these molecules at a membrane would act as a proxy for the magnitude of auxin influx and efflux through the membrane. Cieslak et al. [21] presented several biochemically plausible implementations of this concept, assuming that tally molecules modulate PIN allocation to membrane, thus giving rise to a feedback of auxin fluxes on PIN localization. When a local increase in auxin influx decreases the abundance of PIN proteins in the corresponding part of the membrane, canalization patterns emerge. Conversely, when auxin influx locally increases PIN allocation, convergence points form. The transition between these two regimes is controlled by a change in a single reaction rate. In addition, thanks to its detailed description of the underlying biochemical reactions, the model does not set PIN concentrations at membranes to their steady-state values, and can thus capture potential transient states during fast polarization events. However, it has been tested only on two-dimensional square grids.

In order to explore three-dimensional aspects of auxin patterning during phyllotaxis, we adapt Cieslak’s model to arbitrary irregular 3D tissue geometries, and implement it using the cell complex paradigm. We begin by running the model on a single layer of cells to reproduce convergence point formation. We then investigate how this process is affected by inner cell layers. Finally, we determine under which conditions midveins initiated at convergent points in the epidermal layer can progress in the inner tissue and eventually connect to the already existing vascular system. We show that Cieslak’s mechanism produces dual polarization in three dimensions, and discuss the conditions under which it can happen. Furthermore, in line with previous studies, we highlight the necessity of an additional mechanism for guiding developing veins towards preexisting vasculature.

## Materials and Methods

### Model of biochemical reactions and transport

We adapted to 3D tissues a network of reactions and transport initially formulated by Cieslak and colleagues for two-dimensional grids of square cells (see figure 9 in [21]). The resulting network is represented as a Petri net in Fig 1A (the system of differential equations derived from this net is detailed in S1 Appendix). It is based on the following hypotheses:

**Fig 1.**
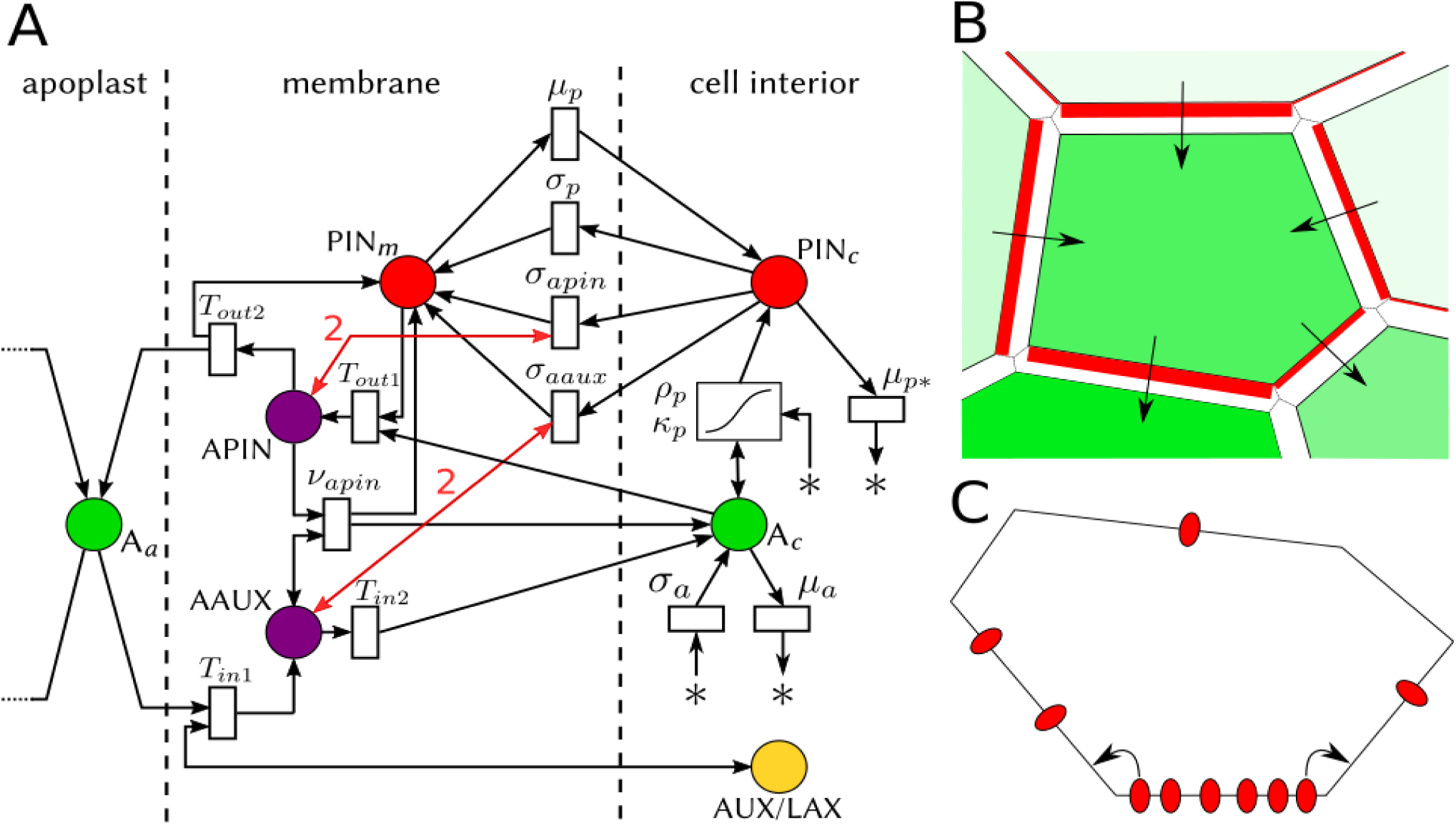
Model description and graphical conventions. (A) Model of cell polarization represented as a Petri net. Chemical species are pictured as circles. A_*a*_: auxin in apoplast; A_*c*_: auxin in cell; PIN_*m*_: membrane-bound PIN proteins; PIN_*c*_: PIN proteins in cell. The symbol * represents reactants or products outside the modeled system (auxin precursors, for instance). Reactions are pictured as rectangles labeled with their rate constants (see main text for an explanation of each term). Arrow orientation indicates whether a species is a reactant or a product in a given reaction. A double-headed arrow indicates a catalytic activity. The two double-headed red arrows represent the impact of the tally molecules APIN and AAUX on PIN exocytosis. They are involved in exocytosis reactions as catalysts with stoichiometric coefficients equal to 2. Other stoichiometric coefficients are equal to 1. An exception is auxin-dependent PIN biosynthesis (rectangle with an S-shaped curve), which follows a more complex kinetics with two parameters (see main text). (B) Graphical convention for simulation results (simplified representation in 2D). Cell interiors are colored in green, with a level of opacity proportional to auxin concentration; for instance the bottom left cell has a much higher concentration than the top cell. Red rectangles on membrane elements picture PIN proteins bound to these elements, with a width proportional to PIN concentration. Arrows make local polarization more apparent; they are deduced from PIN localization. (C) Lateral diffusion of PIN proteins on a cell membrane. Membrane-bound PINs are not fixed at some position on the membrane but freely diffuse on it. This idea is pictured on a 2D section of a cell, with membrane-bound PINs represented as red ellipses. On average, PINs move from regions with higher concentrations to regions with lower concentrations. Since we discretized each cell membrane into a set of polygonal faces (referred to as membrane elements), PIN diffusion was modeled as fluxes between adjacent faces of a given cell membrane, using a discrete Fick’s law.

1) Auxin efflux is a two-step process. In the first step, cellular auxin (A_c_) forms a complex with membrane-bound PIN proteins (PIN_m_):

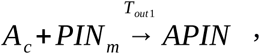

where T_out1_ is the rate of APIN formation. The APIN complex is bound to the membrane. In the second step, APIN spontaneous dissociation releases auxin in the apoplast (A_a_):

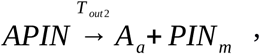

where T_out2_ is the rate of APIN spontaneous dissociation. As a result of this two-step process, the local concentration of APIN is proportional to the local magnitude of auxin efflux (the proportionality constant being the inverse of the rate of APIN dissociation). In this sense, the APIN complex plays the role of a tally molecule measuring auxin efflux.
2) Auxin efflux locally increases exocytosis of cellular PIN (PIN_c_) through the tally molecule APIN:

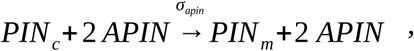

where σ_apin_ is the rate of PIN exocytosis by APIN. Two APIN molecules are needed to transport one PIN from the cytoplasm to the membrane. This ensures quadratic feedback of auxin flux on PIN exocytosis [22-24].
3) Auxin influx also occurs in two steps. First, apoplastic auxin forms a complex with AUX/LAX efflux carriers:

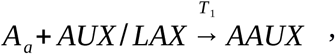

where T_out1_ is the rate of AAUX formation. The AAUX complex is bound to the membrane. In the second step, AAUX spontaneously dissociates:

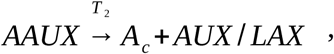

where T_in2_ is the rate of AAUX spontaneous dissociation. After dissociation, auxin is released in the cytoplasm. Paralleling APIN, the local concentration of AAUX is proportional to the local magnitude of auxin influx (the proportionality constant being the inverse of the rate of AAUX dissociation). The AAUX complex plays the role of a tally molecule measuring auxin influx.
4) Auxin influx locally increases PIN exocytosis through the tally molecule AAUX:

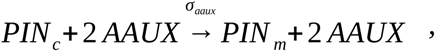

where σ_aaux_ is the rate of PIN exocytosis by APIN. Again, the feedback of auxin flux on PIN exocytosis is quadratic.
5) AAUX catalyzes dissociation of APIN:

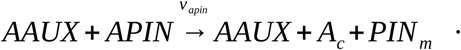 After dissociation of APIN, auxin is released back into the cytosol. ν_apin_ is the rate of APIN dissociation by AAUX and its value depends on auxin concentration in the cell. For low auxin concentration, ν_apin_ is close to zero, so that auxin influx acts only positively on auxin efflux. This leads to convergence point formation. Conversely, for high auxin concentration, ν_apin_ has a high value, so that AAUX rapidly breaks up APIN. Therefore, auxin influx hinders auxin efflux, which leads to canalization. The transition between the two extremal values of ν_apin_ follows a sigmoid function (see S1 Appendix).
6) Besides exocytosis mediated by tally molecules, there are constitutive exo- and endocytosis of PIN proteins, with respective rates σ_p_ and μ_p_.
7) PIN biosynthesis is assumed to be auxin-dependent. The total concentration of PIN in a cell changes according the to formula

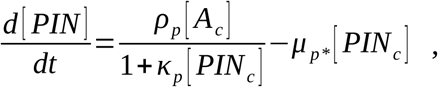

where [A_c_] is the concentration of auxin in the cell, [PIN_c_] is the concentration of cytoplasmic (non-exocytosed) PIN in the cell, ρ^p^ captures the up-regulation of PIN biosynthesis by auxin, κ^p^ controls saturation of PIN biosynthesis, and μ_p*_ is the rate of PIN decay. This is a significant departure from Cieslak’s original scheme, in which a fixed pool of PIN proteins is assumed in every cell. The latter assumption does not held for the full SAM, in which most inner cells contain much less PIN proteins than epidermal cells. Furthermore, auxin-dependent PIN biosynthesis is strongly supported by experiments [25].
8) In each cell, auxin is synthesized and degraded at fixed rates (respectively σ_a_ and μ_a_). In the apoplast, auxin diffuses freely.

### 3D model of tissue

We constructed a 3D model of meristematic tissue. Each tissue template was built in two steps. First, we tessellated a given volume with truncated octohedra. The truncated octohedron has the advantage of filling space, while having a more complex shape than other convex space-filling polyhedra such as cube and prisms, which makes it more realistic as a meristematic cell. In a second step, the vertexes of our mesh were moved by random amounts to introduce irregularities in the template. Each face of the polyhedral cells represents a discrete element of cell membrane. When two cells share a face, this face defines two discrete elements of membrane (one for each cell) and a discrete element of apoplast (shared by both cells).

### Implementation and visualization of simulations

The model was implemented in C++ using the VVe modeling environment, an extension of the VV system [26]. The tissue is represented as a 3D cell complex. The representation and associated topological operations have been adapted from the work of Brisson [27,12]. The simulations are visualized using the following graphical convention (see Fig 1B):

- The opacity of a cell is set proportional to its cellular auxin concentration. A cell devoid of auxin is transparent.
- Cells with a high auxin biosynthesis rate (most of L1 cells) are colored in green.
- Cells with a low auxin biosynthesis rate (inner cells plus a few L1 cells) are colored in white.
- Cells with a high auxin turnover rate (i.e. sink cells) are colored in purple and are fully opaque.
- Membrane-bound PINs on a membrane element are represented as a red plate whose thickness is proportional to PIN concentration.

## Results

### Cieslak’s mechanism can produce auxin convergence points in a single layer of irregular 3D cells

We first run our model on a tissue template consisting of 261 cells arranged in a single planar layer with a roughly circular shape. We assigned a high auxin biosynthesis rate to all cells, except from two subgroups of cells which have a low auxin biosynthesis rate:

- Cells at the border of the template, to account for the fact that tissues surrounding the SAM act more as auxin sinks than as sources.
- The five most central cells, to avoid convergence point formation in this region, in agreement with modeling and experimental data [1,28].

Initial auxin concentration was set to zero in all cells. In the first part of the simulation, auxin concentration increased steadily until instabilities occurred and PINs started to polarize. Then, convergence points formed rapidly (Fig 2A and S1 Movie).

**Fig 2.**
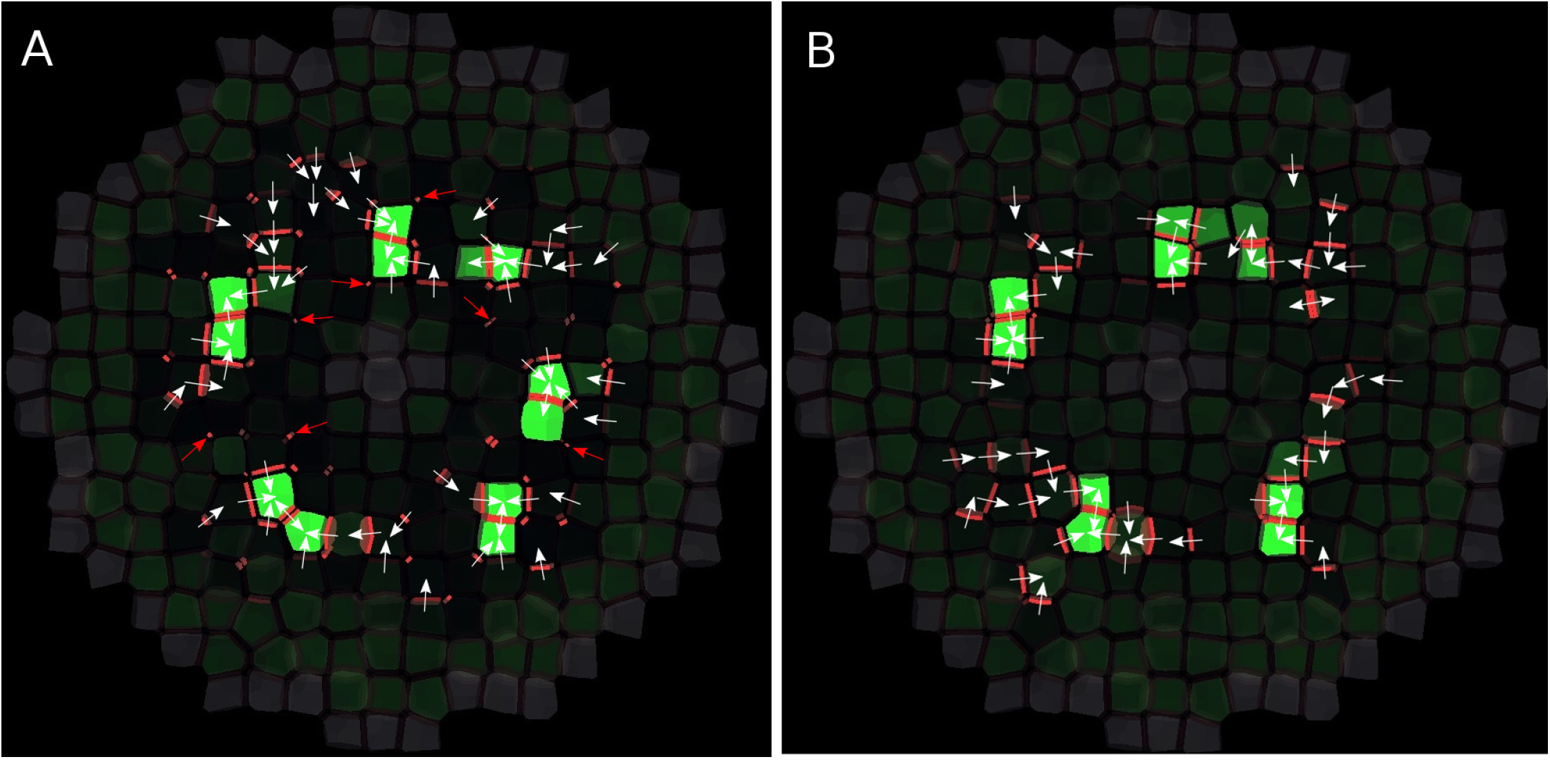
Convergence point formation in a single layer of cells. (A) This image is to be considered as a shoot apical meristem viewed from above. Regions of high cell auxin concentration and PIN polarization appear. White arrows make the local direction of PIN polarization more easily visible. There is a strong tendency for PIN proteins to be allocated on membrane elements with small areas (a few examples are pointed by red arrows). This produces an unrealistic convergence patterns. For explanation of graphical conventions, see Fig. 1B. (B) Same simulation, but with lateral diffusion of membrane-bound PINs. The exaggerated role played by membranes with a small area has disappeared and the pattern of convergence is more realistic, with four convergence points.

A visible unrealistic feature in the convergence pattern obtained from this simulation was that some membrane elements with a small area were favored over elements with a larger area in terms of PIN allocation. PINs tended to accumulate on them. This especially happened when two cells with very different auxin concentrations shared only a very small element of cell wall. Then, an auxin flux tended to establish, which promoted PIN allocation to the corresponding membrane element(s). Since the membrane elements of the two adjacent cells were very small, the flux kept a high magnitude, so equilibrium was reached very slowly. In the meantime, membrane-bound PIN concentration reached unrealistically high values. When the membrane elements involved were extremely small, this could even lead to numerical instabilities.

That aberration has various causes. First, discretizing cell membranes and walls, as it is commonly done in tissue modeling, introduces some distortions in the way fluxes are modeled. This does not represent such a problem in regular (square or hexagonal) cell grids, in which all discrete elements have the same size. Three-dimensionality makes the problem more acute because it spontaneously generates a broader range of areas. Second, it is unrealistic not to set an upper limit to the concentration of membrane-bound PINs since PIN proteins occupy some space on a membrane and thus can not reach too high densities due to steric limitations. A solution could have been to assume that PIN allocation follows some limiting kinetics, for instance Michaelis-Menten. But as this sounded a bit *ad hoc,* we chose instead to assume that membrane-bound PINs diffuse between adjacent membrane elements of the cell, which seemed justified by the fluid nature of the lipidic cell membrane (Fig 1C; see mathematical details in S1 Appendix).

We ran another simulation on the same tissue template, but this time assuming significant lateral diffusion of membrane-bound PIN proteins. We obtained a similar pattern of convergence points, except for the tendency of PINs to accumulate on small membrane elements, which was no longer observed, as expected (Fig 2B and S2 Movie). The PIN lateral diffusion hypothesis did not only eliminate over-accumulation of PINs on small parts of a membrane (since PINs diffused from regions with higher concentrations to adjacent regions with lower concentrations), but it also mitigated the distortions caused by artificial membrane discretization, since the separation between membrane elements was made less strict. Furthermore, it made the model of PIN polarization more dynamic: membrane-bound PINs could then also be reallocated through lateral diffusion, not only through endo- and exocytosis.

### Inner cell layers actively counters leak of auxin from the L1

We considered a tissue template with four layers of cells to address the question of how auxin is kept confined in the L1 until venation begins. We ran several simulations on this template, setting a high auxin biosynthesis rate in L1 cells (except from cells at the border and center, as previously) and a low auxin biosynthesis rate in inner cells (L2, L3, and L4).

We first assumed that only epidermal cells can synthesize PINs, while inner cells are devoid of PINs. Our simulations could not reproduce convergence points in the epidermal layer due to significant leaks of auxin into inner layers. This prevented auxin concentration in the L1 to reach the critical value at which convergence points form. To investigate whether PIN proteins in inner layers could influence auxin patterning in the epidermal layer, we amended our assumption by assigning auxin-dependent PIN biosynthesis to all cells. We could then recover the epidermal patterning (Fig 3 and S3 Movie). This was only possible if the rate of auxin-dependent PIN biosynthesis was high enough, so that PINs were quickly produced in the L2 as soon as auxin was entering L2 cells from the L1. The newly produced PINs polarized towards the L1, obeying up-the-gradient polarization, and successfully countered inward auxin flow until convergence points formed. These results point to an active contribution of L2 PINs to auxin patterning in the epidermal layer. This is in line with observations and simulations by Bayer et al, [8], who also reported upward PIN polarization in the L2. It should be noted, though, that the convergence pattern obtained with four cell layers is a bit blurred compared with the pattern obtained with a single layer.

**Fig 3.**
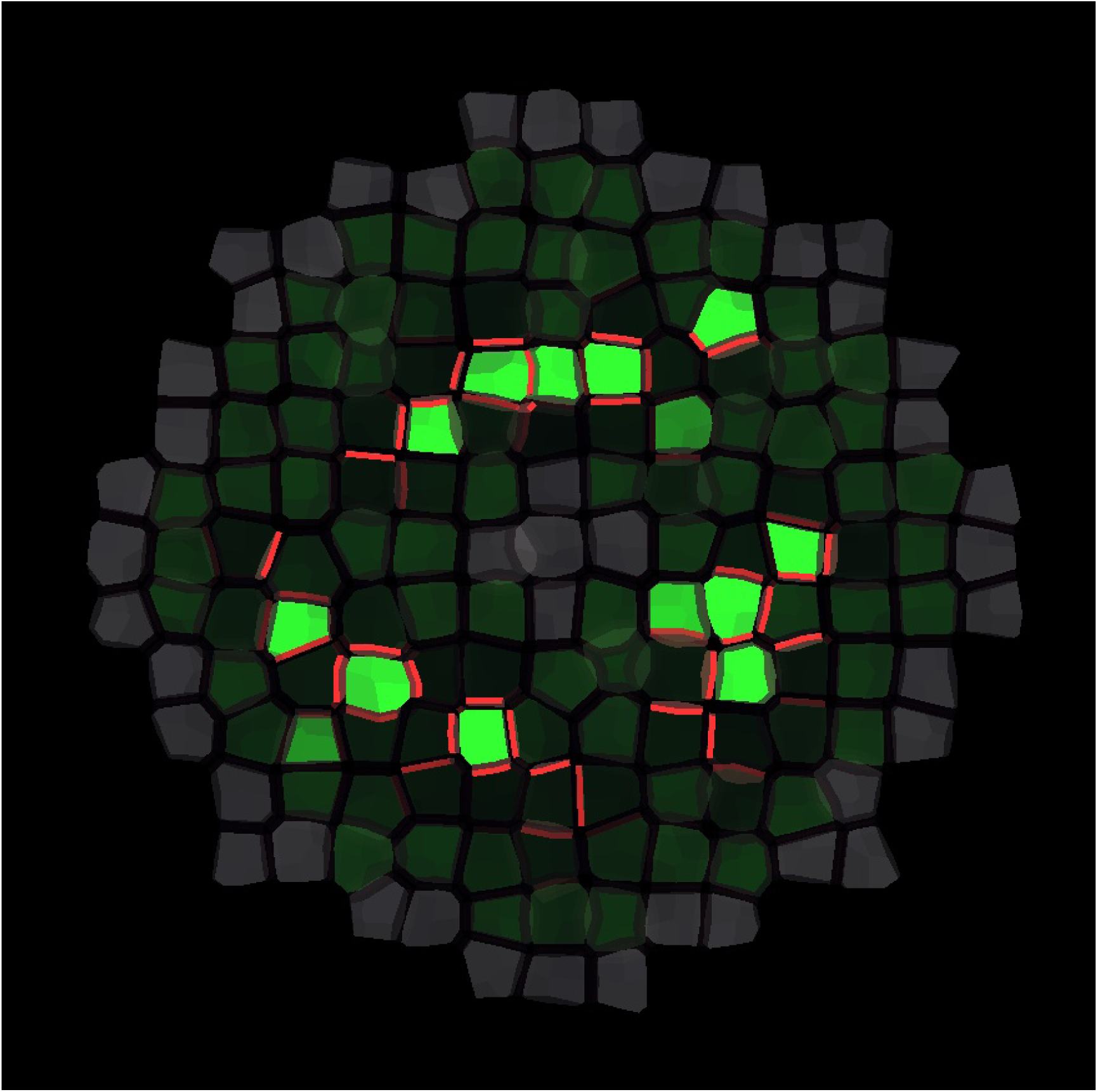
Convergence point formation with four layers and auxin-dependent PIN biosynthesis in all cells. Only the L1 is shown, the inner layers are hidden. Auxin converges towards three groups of cells.

### Convergence points initiate downward veins

Using the same template and parameter values as in the previous simulation, we investigated vein initiation. We found that veins developed from convergence points (Fig 4 and S4 Movie, S5 Movie). Very early in the formation of a convergence point, most PIN proteins were allocated to the bottom membrane of the cells in which auxin converged. Auxin flow was thus locally directed towards the L2 and a midvein was initiated.

**Fig 4.**
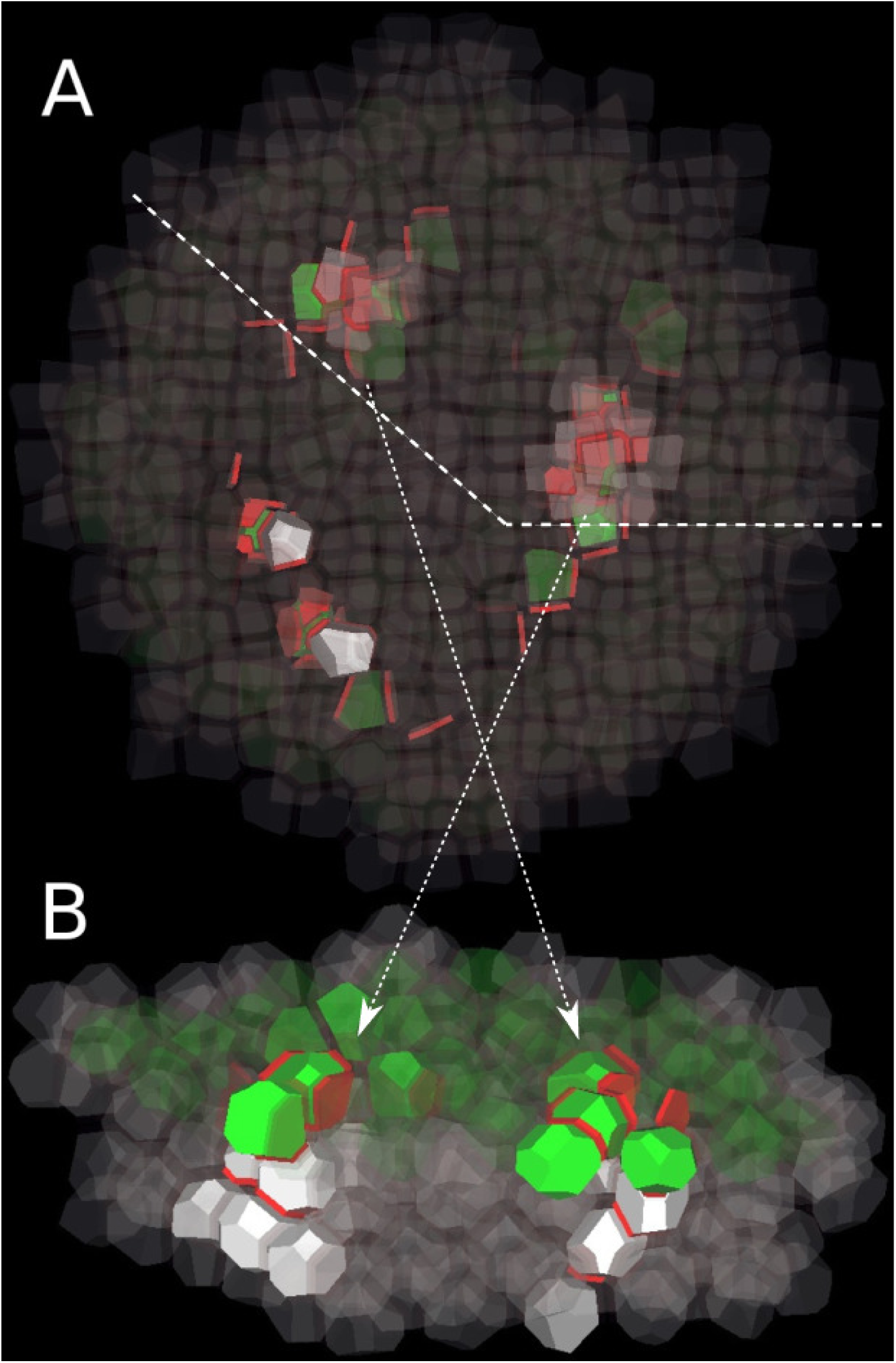
Initiation of veins from convergence points. (A) The tissue is viewed from the bottom, with the bottom cell layer L4 in the foreground and the L1 in the background (visible through the transparency of the cells). Cells in inner layers are white, with opacity proportional to their auxin concentration. Three convergence points in the L1 are visible as groups of green cells. From these convergence points, auxin is polarly transported downward into inner tissue. (B) On a longitudinal cutaway following the dashed line in (A), vein initiation appears more clearly. The arrows show the correspondence between both panels.

It turned out that the initiation of veins was possible only if auxin importer concentration in inner cell layers was at least as high as in the L1. This is in contrast to experimental results by Kierzkowski et al. [7] and Bainbridge et al. [29], who did not observe auxin influx carrier AUX1 nor LAX1 outside the L1. This suggests the existence of other auxin importers than AUX1 and LAX1 in the inner tissue.

### Developing veins are only weakly attracted by auxin sinks

In order to investigate how veins are progressing in the inner tissue and potentially connect to the pre-existing vasculature, we modified the previous template by assigning three evenly-spaced cells in the bottom layer (L4) as sinks. Sink cells had a high auxin turnover rate, which was intended to mimic the effect of a functional vein draining auxin. Thus, each sink cell induced a local gradient of auxin concentration centered around itself.

In our simulations, when a cell was exporting auxin towards a neighboring cell in the inner tissue, the auxin concentration in the latter cell was increasing until it started to polarize in turn towards a third cell, and so on. Veins developed this way, from convergence points towards regions with lower auxin concentration. Therefore, they progressed globally downward, away from L1 auxin-producing cells (Fig 5 and S6 Movie, S7 Movie, S8 Movie). Often, a vein did not develop as a single sequence of cells, but instead as a bundle of such parallel sequences. Here, lateral diffusion of PIN proteins makes a clear difference. Without lateral diffusion of PINs, each vein formed as a single cell sequence, and this sequence followed preferentially paths connecting cells through very small apoplast element. This was the same effect we had seen in convergence point formation in the epidermal layer, where small membranes were favored over larger ones. Again, this did not fit observations, so lateral diffusion of PINs appears as a necessary feature for modeling venation in a 3D irregular tissue.

**Fig 5.**
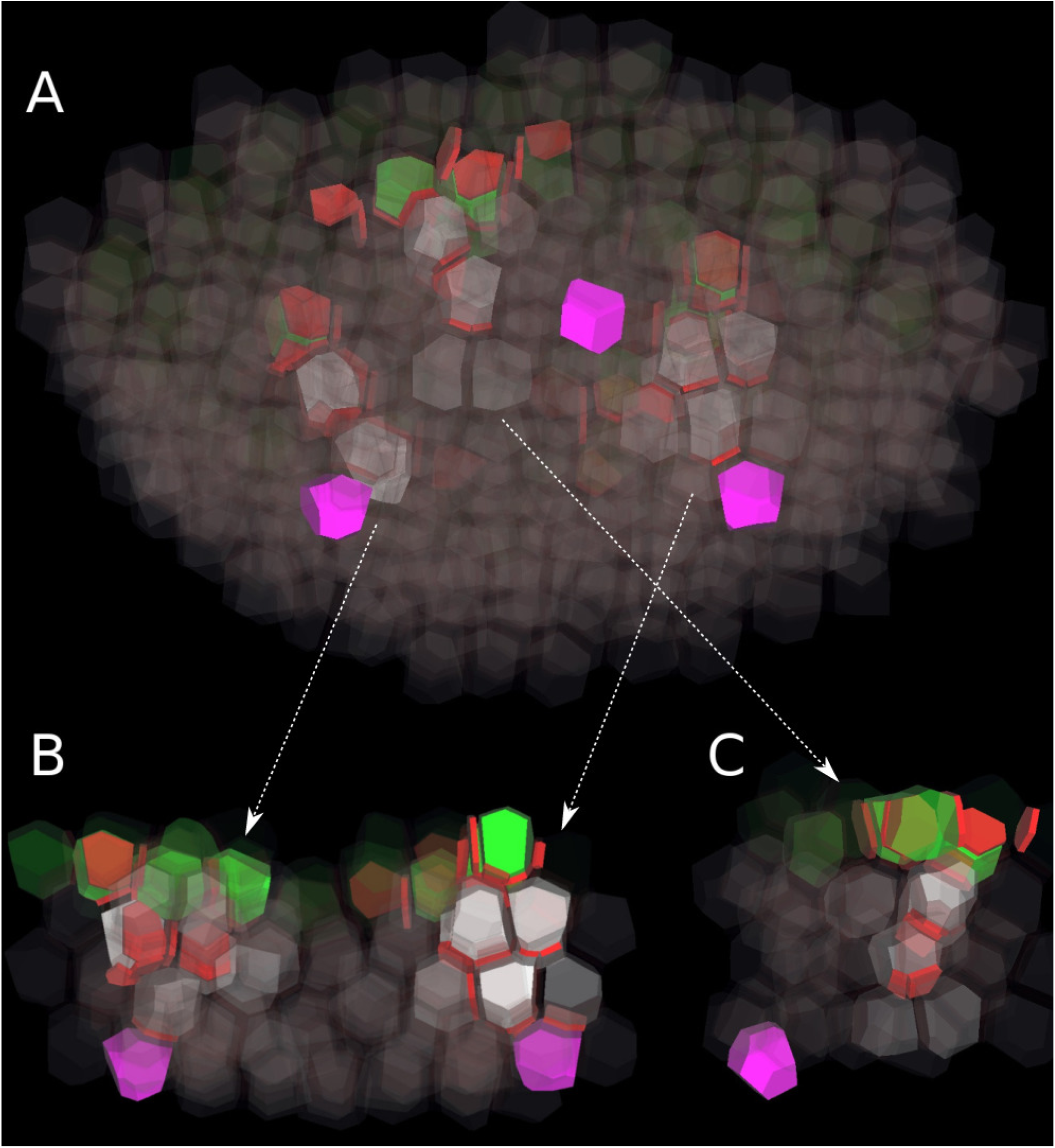
Progression of initiating veins in inner layers. (A) The tissue is viewed obliquely from the bottom. The three sink cells (in purple) are located at different positions in the deepest layer (L4). In this simulation, auxin forms convergence points in the L1 (green cells), and then is canalized into the inner tissue (white cells). Two veins (on left and right) eventually connect to a sink, while a third one (middle) gets lost in the tissue. (B) Longitudinal cutaway with the two veins which connect to a sink. (C) Longitudinal cutaway with the vein which fails to reach a sink. The arrows show the correspondence between the oblique view and the cutaways.

Since developing veins globally progressed towards lower auxin concentrations, and since sink cells laid at the center of low auxin concentration regions, it was expected that all veins were going to eventually reach a sink. However, local auxin gradients surrounding sinks were very shallow and only had a short-range attraction power on developing veins. Veins which did not pass close to a sink missed their target and got lost in the tissue (Fig 5C and S8 Movie). Therefore, although Cieslak’s mechanism does reproduce several features of vein formation in a 3D tissue, it does not provide an efficient way to find sinks.

### Assuming a vein attraction factor (VAF) explains connection of developing veins to already existing vasculature

The poor capacity of developing veins to find sinks in our simulation is suggestive of an additional mechanism for guiding veins towards the already existing vasculature. Bayer et al. [8] reached a similar conclusion. This mechanism could take the form of facilitated diffusion of auxin through plasmodesmata, which has been proven by Smith and Bayer [30] to reliably and robustly connect sources to sinks. Both facilitated diffusion and polar transport could operate in parallel, with facilitated diffusion first defining a preferred route to the closest sink, and polar auxin transport then establishing polarization along this route. But there is little experimental support for facilitated diffusion of auxin in plant tissue. Therefore, following Bayer et al. [8], we resorted to an alternative model which posits a vein attraction factor (VAF) emitted by existing vasculature.

In the original model by Bayer et al. [8], the vein attraction factor diffuses from cell to cell. Every cell can measure VAF concentrations in its neighbors and biases PIN allocation towards the membrane elements facing the cells with highest VAF concentrations. In Cieslak’s model, however, there is no such remote sensing between cells. Its biochemical plausibility rests on the fact that all reactions occurs locally, between a cytoplasm and a membrane, or between a membrane and neighboring apoplast. We propose an alternative VAF behavior, which is compatible with the locality of Cieslak’s model. In our scheme, sink cells release VAF in their neighboring apoplast at a fixed rate. The VAF diffuses within the apoplast continuum, never reentering cells. However, it can bind to membranes and unbind from them, at some fixed rates. The concentration of bound VAF on a membrane element favors PIN exocytosis toward this membrane element.

We implemented this mechanism (see details in S1 Appendix) and found in our simulations that VAF gradients could establish and set a local polarity around every sink cell. The spatial range of VAF-induced polarity depends on VAF production rate and diffusion coefficient. If these values were adequately set, every initiating vein eventually headed and connected to the nearest sink cell (Fig 6 and S9 Movie, S10 Movie, S11 Movie). The VAF hypothesis introduced an additional feature to vein development. Although the cells surrounding a sink had, at the beginning, low auxin and PIN contents, they rapidly polarized towards their neighbor sink. As time went by and VAF diffused farther away, second-order neighbor cells started to polarize towards the sink. When an initiating vein reached a cell already polarized towards a sink, this polarization got reinforced thanks to the high influx of auxin from the vein and the PIN production induced. PIN polarization was then firmly established between the source and the sink. To sum up, the complete vein was the result of the encounter of two opposed movements: the progression of an initiating vein towards a sink, and the expansion of a polarized region centered around this sink. In the initiating vein, cells had high auxin concentrations since each cell had to reach a threshold concentration before it could polarize towards another cell and made the vein progress. But, when the initiating vein reached the polarized region near the sink, auxin could flow rapidly to the sink, without accumulating. Thus, at the moment the polarized path was established, cells in the upper part of the vein had high auxin concentrations while cells closer to the sinks had low concentrations. The former cells kept their high concentration whereas the latter cells only very slowly increased their concentration.

**Fig 6.**
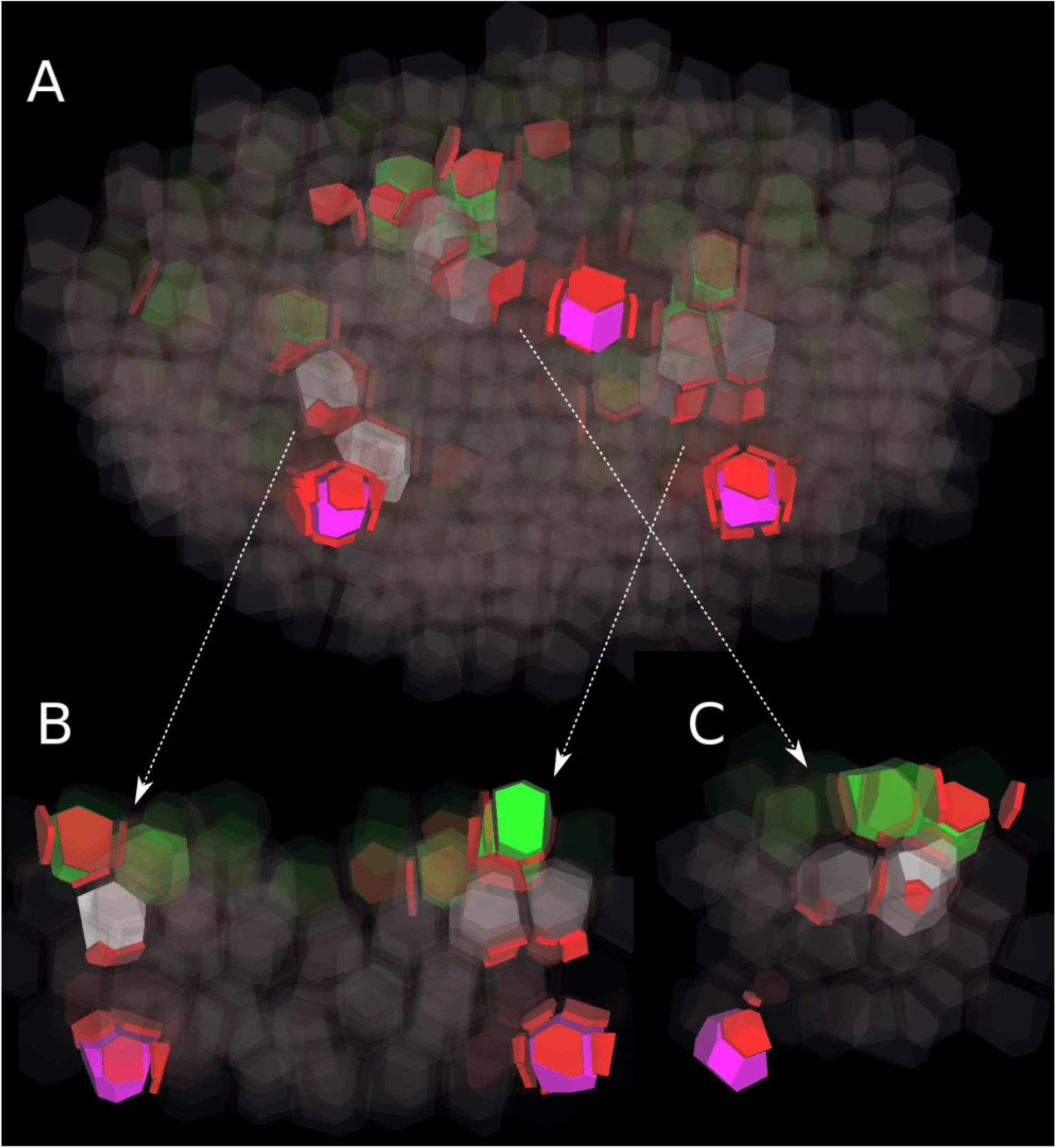
Vein progression with Vascular Attraction Factor. Simulation performed on the same tissue template as in figure 5, with the same spatial distribution of auxin production and turnover, and same color code. The only additional ingredient is the production of a Vascular Attraction Factor (VAF), produced by sink cells and biasing vein progression towards higher concentration of VAF. As a result, all veins eventually connect to a sink. (A) The tissue is viewed obliquely from the bottom. (B) Longitudinal cutaway with two veins (arrows indicate correspondence with panel A. (C) Longitudinal cutaway with the third vein. Note that auxin has no longer time to accumulate in the vein portions near sinks, since connection to sink occurs very early.

## Discussion

The shoot apical meristem is a challenging system in our understanding of auxin patterning due to the overlap in time and space of different auxin transport regimes in an irreducible three-dimensional geometry. It poses both a conceptual and a technical problem. From a conceptual point of view, the co-localization of up-the-gradient and with-the-flux polarizations in the same cells, with the same efflux carriers, strongly suggests a common mechanism, while most models so far have addressed each polarization regime separately with incompatible hypotheses. From a technical point of view, building a 3D computational model of plant tissue including both symplast and apoplast, plus polar auxin transport, is a challenge which had never been taken up. As a significant first step towards that objective, we built a model of simplified meristematic tissue on which a biochemically plausible mechanism of auxin transport was implemented. This approach reproduced key features of 3D phyllotactic patterning and offered new insights in the dynamics of PIN1 polarization in the shoot apical meristems.

We emphasize the importance of membrane-bound PIN diffusion in auxin patterning, for both convergence point and canalization. This has been overlooked in 2D models, in which the sizes of the discrete membrane elements are approximately of the same order of magnitude, and usually quite large. But in 3D geometry, the higher complexity of the topology results in a very wide range of areas for discrete membrane elements, with occasionally extremely small values. This compartmentalization becomes then overly artificial and ignores the intrinsic continuity of the lipid cell membrane in which bound PIN proteins can move laterally. It leads to aberrant accumulation of PINs on a few very tiny membrane elements. Assuming lateral diffusion of PIN proteins partially overcomes this issue by smoothing the barriers between neighbor discretized elements of the same cell membrane. It should be noted, however, that membrane-bound PINs display a more complex behavior, due to mechanisms limiting their lateral movements [31]. They tend to form clusters in which lateral mobility is strongly reduced. Only a minor fraction of membrane-bound PINs are unclustered and laterally mobile [32]. Future models could investigate in more details the significance of PIN clustering for cell polarization at tissue scale.

We demonstrated that inner cell layers, especially the L2, significantly contribute to auxin patterning in the L1. The higher auxin biosynthesis rate in the epidermal layer is not sufficient by itself to reach the critical concentration required for the formation of convergence points. In the early stages of phyllotactic patterning, auxin has to be kept confined in the L1. We showed that this confinement can be performed by up-the-gradient polarization in the L2, where PIN proteins polarize upward, towards auxin-rich L1 cells. A necessary condition is that auxin-dependent PIN production is fast enough to react quickly to the first auxin leaks into the L2. Bayer et al. [8] assumed reduced symplasmic communication between L1 and inner tissues, based on experimental evidence [33]. We did not make the assumption and found that PIN polarization alone can efficiently confine auxin in the L1 until midvein initiation. We also found that mutual influences between cell layers have a slight but non-negligible impact on the final pattern. This suggests that precise and realistic descriptions of surface patterning should take into account effects from the underlying tissue.

It turned out that midveins cannot develop if the concentration of auxin influx carriers is lower in the inner tissue than in the L1. This seemingly contradicts experimental results [7,29] showing that auxin influx carriers AUX1 and LAX1 are almost exclusively present in the epidermal layer. But at the same time, Bainbridge et al. [29] reported the expression of another auxin importer, LAX2, in the initiating vein. LAX2 has also been shown to regulate vascular patterning in cotyledons [34]. Our model does not exclude the possibility that auxin influx carriers present in the inner tissue could be restricted to the cells forming the initiating veins. However, in this case, they have to be expressed from the very beginning of vein initiation to make it possible. Since the mechanism inducing LAX2 expression is not known, amending the model in this direction seems both speculative and premature.

Our investigation also pointed out that auxin gradients induced by the existing vascular system are too shallow to efficiently attract initiating veins. This fact had been already noted in 2D simulations performed by Bayer et al. [8], and is linked to the more general problem of establishing polarity within a tissue. Various models of tissue polarity have been developed (see a comparative study in [18]). In the shoot apical meristem of Arabidopsis, the region in which a polarity has to be established to guide an initiating vein is relatively small. Therefore, a simple mechanism based on a putative diffusive molecule is sufficient, as already outlined by Bayer et al. [8]. We proposed a detailed process for the action of such a Vein Attraction Factor. We assumed that the VAF is transported apoplastically, which seemed to us more realistic than the symplastic transport first proposed by Bayer et al. [8].

This vein attraction mechanism raises new questions about source-sink connections. Classically, an initiating vein is viewed as progressing in a tissue from a source until it reaches a sink. In our simulations, each sink builds up an expanding polarity field centered around itself. When the progressing vein meets the expanding polarity field, the connection is almost fully established. Then, the vein progresses very quickly and directly to the sink, locally increasing polarity in its wake. The relative contribution of each process---free progression of the initiating vein and expansion of the polarity field---depends on the biosynthesis rate and diffusion coefficient of the VAF, but also on the precise timing of vein initiation and VAF release. Further experimental studies are needed to improve our understanding of how primordia connect to the vascular system. Modeling efforts are hindered by the unknown nature of the putative VAF.

Another obvious limitation to our model is that it does not reproduce phyllotactic patterns. This is essentially because it does not include tissue growth, which plays a crucial role in the dynamics of phyllotactic patterning. However, simulating meristem growth with mechanical processes would add another layer of technical complexity and requires much more development and computational resources. Making the size and shape of the template closer to an actual meristem is also a necessary improvement for future models to gain more realism.

It can also be argued that the PIN polarization mechanism used in our model is questionable since there is no evidence for auxin complexes. Yet, the core of the mechanism is more generic than it looks at first sight. Cieslak et al. [21] proposed various other implementations of it, reflecting different biochemical assumptions. Since the precise reactions underlying the feedback of auxin transport on PIN localization are unknown, it is difficult to discriminate between these implementations. Other ones can be designed based on new experimental findings. For instance, the fact that convergence point formation and vein initiation seem to be associated with two distinct groups of auxin influx carriers could be exploited to amend the current reaction network. Alternatives will probably emerge in the near future from experimental studies, and 3D models like ours will be valuable tools to thoroughly assess their explanatory powers in terms of auxin pattern formation.

## Acknowledgments

We thank Richard S. Smith for sharing valuable ideas and for useful comments on the manuscript. FPH thanks Agata Burian, Sarah J. Robinson, and Hagen Reinhardt for helpful discussions.

## Supporting information

**S1 Appendix. Technical details on models used.**

**S1 Movie. Convergence point formation in a single layer of cells, without lateral diffusion of PINs.**

**S2 Movie. Convergence point formation in a single layer of cells, with lateral diffusion of PINs.**

**S3 Movie. Convergence point formation with four layers and auxin-dependent PIN biosynthesis in all cells.** *Only the L1 is shown, the inner layers are hidden. Auxin converges towards three groups of cells.*

**S4 Movie. Initiation of veins from convergence points.** The tissue is viewed from the bottom, with the bottom cell layer L4 in the foreground and the L1 in the background (visible through the transparency of the cells). Cells in inner layers are white, with opacity proportional to their auxin concentration. Three convergence points in the L1 are visible as groups of green cells. From these convergence points, auxin is polarly transported downward into inner tissue.

**S5 Movie. Initiation of veins from convergence points, viewed on a longitudinal cutaway.** The tissue is viewed from the side, on a longitudinal cutaway, like in Fig 4.

**S6 Movie. Progression of initiating veins in inner layers.** The tissue is viewed obliquely from the bottom, like in Fig 5A. The three sink cells (in purple) are located at different positions in the deepest layer (L4).

**S7 Movie. Progression of two initiating veins toward sinks.** The tissue is viewed from the side, on a longitudinal cutaway showing two veins, like in Fig 5B.

**S8 Movie. Progression of an initiating vein failing to reach a sink.** The tissue is viewed from the side, on a longitudinal cutaway showing one vein, like in Fig 5C.

**S9 Movie. Progression of initiating veins in inner layers, with VAF.** The tissue is viewed obliquely from the bottom, like in Fig 6A.

**S10 Movie. Progression of two initiating veins toward sinks, with VAF.** The tissue is viewed from the side, on a longitudinal cutaway showing two veins, like in Fig 6B.

**S11 Movie. Progression of an initiating vein toward a sink, with VAF.** The tissue is viewed from the side, on a longitudinal cutaway showing one vein, like in Fig 6C.

## References

1. Barbier de Reuille P, Bohn-Courseau I, Ljung K, Morin H, Carraro N, Godin C, et al. Computer simulations reveal properties of the cell-cell signaling network at the shoot apex in Arabidopsis. PNAS. 2006;103(5):1627–1632. doi:10.1073/pnas.0510130103.

2. Jönsson H, Heisler MG, Shapiro BE, Meyerowitz EM, Mjolsness E. An auxin-driven polarized transport model for phyllotaxis. Proc Natl Acad Sci U S A. 2006;103(5):1633–8. doi:10.1073/pnas.0509839103.

3. Smith RS, Guyomarc’h S, Mandel T, Reinhardt D, Kuhlemeier C, Prusinkiewicz P. A plausible model of phyllotaxis. Proc Natl Acad Sci U S A. 2006;103(5):1301–6. doi:10.1073/pnas.0510457103.

4. Stoma S, Lucas M, Chopard J, Schaedel M, Traas J, Godin C. Flux-based transport enhancement as a plausible unifying mechanism for auxin transport in meristem development. PLoS Comput Biol. 2008;4(10):e1000207. doi:10.1371/journal.pcbi.1000207.

5. Reinhardt D, Pesce ER, Stieger P, Mandel T, Baltensperger K, Bennett M, et al. Regulation of phyllotaxis by polar auxin transport. Nature. 2003;426(6964):255–260. doi:10.1038/nature02081.

6. Banasiak A. Putative dual pathway of auxin transport in organogenesis of Arabidopsis. Planta. 2011;233(1):49–61. doi:10.1007/s00425-010-1280-0.

7. Kierzkowski D, Lenhard M, Smith R, Kuhlemeier C. Interaction between meristem tissue layers controls phyllotaxis. DevCell. 2013;26(6):616–628. doi:10.1016/j.devcel.2013.08.017.

8. Bayer EM, Smith RS, Mandel T, Nakayama N, Sauer M, Prusinkiewicz P, et al. Integration of transport-based models for phyllotaxis and midvein formation. Genes & Development. 2009;23(3):373–384. doi:10.1101/gad.497009.

9. Runions A, Smith RS, Prusinkiewicz P. In: Zažıımalová E, Petrášek J, Benková E, editors. Computational Models of Auxin-Driven Development. Vienna: Springer Vienna; 2014. p. 315–357.

10. Gruel J, Landrein B, Tarr P, Schuster C, Refahi Y, Sampathkumar A, et al. An epidermis-driven mechanism positions and scales stem cell niches in plants. Science Advances. 2016;2(1). doi:10.1126/sciadv.1500989.

11. Prusinkiewicz P, Lane B. In: Capasso V, Gromov M, Harel-Bellan A, Morozova N, Pritchard LL, editors. Modeling Morphogenesis in Multicellular Structures with Cell Complexes and L-systems. Berlin, Heidelberg: Springer Berlin Heidelberg; 2013. p. 137–151.

12. Yoshida S, de Reuille PB, Lane B, Bassel GW, Prusinkiewicz P, Smith RS, et al. Genetic Control of Plant Development by Overriding a Geometric Division Rule. Developmental Cell. 2014;29(1):75–87. doi:10.1016/j.devcel.2014.02.002.

13. Merks RMH, de Peer YV, Inzé D, Beemster GTS. Canalization without flux sensors: a traveling-wave hypothesis. Trends in Plant Science. 2007;12(9):384–390. doi:10.1016/j.tplants.2007.08.004.

14. Bennett T, Hines G, Leyser O. Canalization: what the flux? Trends in Genetics. 2014;30(2):41–48. doi:10.1016/j.tig.2013.11.001.

15. O’Connor DL, Runions A, Sluis A, Bragg J, Vogel JP, Prusinkiewicz P, et al. A Division in PIN-Mediated Auxin Patterning during Organ Initiation in Grasses. PLoS Comput Biol. 2014;10(1):e1003447. doi:10.1371/journal.pcbi.1003447.

16. Wabnik K, Kleine-Vehn J, Balla J, Sauer M, Naramoto S, Reinöhl V, et al. Emergence of tissue polarization from synergy of intracellular and extracellular auxin signaling. Molecular Systems Biology. 2010;6(1). doi:10.1038/msb.2010.103.

17. Abley K, De Reuille PB, Strutt D, Bangham A, Prusinkiewicz P, Marée AFM, et al. An intracellular partitioning-based framework for tissue cell polarity in plants and animals. Development. 2013;140(10):2061–2074. doi:10.1242/dev.062984.

18. Abley K, Sauret-Güeto S, Marée AF, Coen E. Formation of polarity convergences underlying shoot outgrowths. eLife. 2016;5:e18165. doi:10.7554/eLife.18165.

19. Heisler MG, Jönsson H. Modeling Auxin Transport and Plant Development. Journal of Plant Growth Regulation. 2006;25(4):302–312. doi:10.1007/s00344-006-0066-x.

20. Coen E, Rolland-Lagan AG, Matthews M, Bangham JA, Prusinkiewicz P. The genetics of geometry. PNAS. 2004;101(14):4728–4735. doi:10.1073/pnas.0306308101.

21. Cieslak M, Runions A, Prusinkiewicz P. Auxin-driven patterning with unidirectional fluxes. J Exp Bot. 2015;66(16):5083–5102. doi:10.1093/jxb/erv262.

22. Mitchison GJ. A model for vein formation in higher plants. Proceedings of the Royal Society of London B: Biological Sciences. 1980;207(1166):79–109. doi:10.1098/rspb.1980.0015.

23. Mitchison GJ. The polar transport of auxin and vein patterns in plants. Philosophical Transactions of the Royal Society of London B: Biological Sciences. 1981;295(1078):461–471. doi:10.1098/rstb.1981.0154.

24. Rolland-Lagan AG, Prusinkiewicz P. Reviewing models of auxin canalization in the context of leaf vein pattern formation in Arabidopsis. The Plant Journal. 2005;44(5):854–865. doi:10.1111/j.1365-313X.2005.02581.x.

25. Vieten A, Vanneste S, Wisniewska J, Benková E, Benjamins R, Beeckman T, et al. Functional redundancy of PIN proteins is accompanied by auxin-dependent crossregulation of PIN expression. Development (Cambridge, England). 2005;132:4521–4531. doi:10.1242/dev.02027.

26. Smith C, Prusinkiewicz P, Samavati F. In: Pfaltz JL, Nagl M, Böhlen B, editors. Local Specification of Surface Subdivision Algorithms. Berlin, Heidelberg: Springer Berlin Heidelberg; 2004. p. 313–327.

27. Brisson E. Representing geometric structures ind dimensions: Topology and order. Discrete & Computational Geometry. 1993;9(4):387–426. doi:10.1007/BF02189330.

28. Vernoux T, Brunoud G, Farcot E, Morin V, Van den Daele H, Legrand J, et al. The auxin signalling network translates dynamic input into robust patterning at the shoot apex. Molecular systems biology. 2011;7:508. doi:10.1038/msb.2011.39.

29. Bainbridge K, Guyomarc’h S, Bayer E, Swarup R, Bennett M, Mandel T, et al. Auxin influx carriers stabilize phyllotactic patterning. Genes & development. 2008;22:810–823. doi:10.1101/gad.462608.

30. Smith RS, Bayer EM. Auxin transport-feedback models of patterning in plants. Plant Cell Environ. 2009;32(9):1258–1271. doi:10.1111/j.1365-3040.2009.01997.x.

31. Martinière A, Lavagi I, Nageswaran G, Rolfe DJ, Maneta-Peyret L, Luu DT, et al. Cell wall constrains lateral diffusion of plant plasma-membrane proteins. Proceedings of the National Academy of Sciences of the United States of America. 2012;109:12805–12810. doi:10.1073/pnas.1202040109.

32. Kleine-Vehn J, Wabnik K, Martinière A, Langowski L, Willig K, Naramoto S, et al. Recycling, clustering, and endocytosis jointly maintain PIN auxin carrier polarity at the plasma membrane. Molecular systems biology. 2011;7:540. doi:10.1038/msb.2011.72.

33. Rinne PL, van der Schoot C. Symplasmic fields in the tunica of the shoot apical meristem coordinate morphogenetic events. Development. 1998;125(8):1477–1485.

34. Péret B, Swarup K, Ferguson A, Seth M, Yang Y, Dhondt S, et al. AUX/LAX genes encode a family of auxin influx transporters that perform distinct functions during Arabidopsis development. The Plant cell. 2012;24:2874–2885. doi:10.1105/tpc.112.097766.

